# High-throughput microfluidic characterization of erythrocyte shapes and mechanical variability

**DOI:** 10.1101/488189

**Authors:** Felix Reichel, Johannes Mauer, Ahmad Ahsan Nawaz, Gerhard Gompper, Jochen Guck, Dmitry A. Fedosov

## Abstract

The motion of red blood cells (RBCs) in microchannels is important for microvascular blood flow and biomedical applications such as blood analysis in microfluidics. The current understanding of the complexity of RBC shapes and dynamics in microchannels is mainly based on several simulation studies, but there are a few systematic experimental investigations. Here, we present a combined study, which systematically characterizes RBC behavior for a wide range of flow rates and channel sizes. Even though simulations and experiments generally show good agreement, experimental observations demonstrate that there is no single well-defined RBC state for fixed flow conditions, but rather a broad distribution of states. This result can be attributed to the inherent variability in RBC mechanical properties, which is confirmed by a model that takes the variation in RBC shear elasticity into account. This represents a significant step toward a quantitative connection between RBC behavior in microfluidic devices and their mechanical properties, which is essential for a high-throughput characterization of diseased cells.

**STATEMENT OF SIGNIFICANCE:** The ability to change shape is crucial for the proper functioning of red blood cells under harsh conditions in the microvasculature, since their shapes strongly affect the flow behavior of whole blood. Our results from simulations and systematic experiments reveal the shapes and dynamics of red blood cells for different flow conditions and channel dimensions, generally in good agreement. However, in the experiments, cells do not exhibit a single well-defined shape for fixed flow conditions. We show that this distribution of shapes can be attributed to the variability in mechanical properties of red blood cells.

## INTRODUCTION

During their 120-day lifespan, red blood cells (RBCs) withstand tremendous stresses and deformations as they travel hundreds of kilometers through the vascular system. Their distinctive biconcave discocyte shape has evolved to meet the requirements for large, repetitive and reversible deformations. The ability to deform and change shape is vital for RBCs, as they pass vessels whose diameter is smaller than their own size (1, 2). Changes in elastic properties and shape of RBCs are of great interest in microcirculation research, because they affect many essential microvascular processes, including blood flow resistance (3–5), the viscosity of whole blood (6–9), ATP release and oxygen delivery (10–12). Alterations in RBC material properties and shape are often associated with different blood diseases and disorders (13, 14), such as sickle cell anemia (15, 16), diabetes mellitus (17, 18), and malaria (19–21). The importance of RBC behavior in flow also motivates large research efforts in blood processing and analysis using microfluidic devices (22–26).

Stimulated by the interest in RBC behavior in microvasculature and microfluidics for blood analysis, a number of experiments and simulations with RBCs flowing in narrow channels and capillaries have been performed (3, 4, 27–33). Typical deformed shapes are so-called parachutes and slippers (4, 5, 34). Parachutes are characterized by a parachute-like shape and a position in the channel center. Slippers take an off-center position in flow and exhibit a tank-treading motion of the RBC membrane due to shear gradients. The occurrence of different RBC shapes is governed by several parameters, including flow rate, cell confinement characterized by the ratio between cell diameter and channel size, RBC elastic properties (e.g. shear elasticity, bending rigidity), and the viscosity ratio *λ* between RBC cytosol and suspending medium (or plasma) (4, 5, 34). Recent simulation studies in two dimensions (2D) (5, 34) and three dimensions (3D) (4) predict a RBC state diagram with various shapes and dynamics for a wide range of flow rates and cell confinements. Even though 2D simulations qualitatively capture some of the RBC states, it is clear that the motion of RBCs is inherently three dimensional and the correspondence between 2D and 3D diagrams is at best qualitative. For example, the 3D simulations (4) predict a tumbling state, where a RBC flips like a coin in Poiseuille flow within a very narrow channel, which does not exist in 2D and has not been observed experimentally so far. One of the limitations of this simulation study (4) is that it was performed for the viscosity ratio *λ* = 1, while under physiological conditions, *λ* ≈ 5 (35). Furthermore, there exists no systematic experimental investigation, which would corroborate or challenge the simulated state diagram of RBC shapes and dynamics in microchannels, even though several RBC states (e.g. parachutes, slippers) have been observed experimentally. Moreover, recent simulations of a RBC in rectangular channels (33) suggest that there might be bistable states even for fixed flow conditions, depending on the initial position of a RBC as it enters the channel, which might indicate a sensitive dependence on the channel geometry. These gaps in our fundamental understanding of RBC behavior in microchannels prevent a knowledge-based design of efficient microfluidic devices for blood analysis.

In this work, we present a combined experimental and simulation study of RBC behavior in microchannels in order to address the challenges above. Microfluidic experiments are performed for a wide range of flow conditions, covering a wide range of cell velocities (between 0.1 mm/s and 4 mm/s) and channel sizes (between 9.75 *µ*m and 14.75 *µ*m in hydraulic diameter). This constitutes the data base for the analysis of about 35 000 cells and allows, for the first time, a systematic construction of an experimental RBC state diagram. Simulations are performed for *λ* ≈ 5 and are in good agreement with experimental observations. A remarkable experimental result is that, for fixed flow conditions, there is no single well-defined RBC state, but rather a distribution of shapes, while simulations generally show a single RBC state. This outcome arises from the inherent variability in RBC mechanical properties, which has to be taken into account. Empirical approximation of the inherent variation in RBC shear elasticity leads to RBC state distributions, which agree well with experimental observations. Furthermore, the experiments show the existence of tumbling polylobe shapes in large enough channels at high flow rates, which were before only observed in pure shear flow (9, 36). The combination of experimental and simulation results allows us to fill several gaps in understanding of RBC behavior in microchannels and to make a step toward a quantitative characterization of RBC mechanical properties and their variability. Future microfluidic investigations will reveal how diseases affect cell mechanical properties and can be used for a precise blood analysis. Such studies will also advance our understanding of the effect of different RBC shapes and dynamics on microvascular blood flow in health and disease.

## MATERIALS AND METHODS

### RBC model

RBC membrane is modeled by a triangulated network of springs, which incorporates stretching, bending, and area-compression resistance (37, 38). The membrane is made of *N* = 3000 vertices. The model also contains area-and volume-conservation constraints to mimic incompressibility of the lipid bilayer and cell’s cytosol. The RBC model employs a stress-free shape of the elastic spring network, corresponding to a spheroidal shape with a reduced volume of 0.96, because recent simulation studies (36, 39, 40) suggest that a nearly spherical stress-free shape best reproduces experimental results for the tumbling-to-tank-treading transition at low viscosity contrasts (41, 42). Alternatively, a stress-free condition can also be incorporated into the bending potential through an inhomogeneous spontaneous curvature (43). RBC size is characterized by the effective RBC diameter 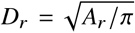,where *A*_*r*_ is the RBC membrane area. Table 1 summarizes the parameters for the RBC model in units of *D*_*r*_ and the thermal energy *k*_*B*_*T*, and the corresponding average values for a healthy RBC in physical units. To relate simulation parameters to the physical properties of RBCs, we need a basic length and energy scales, while a relation for time scale is based on the characteristic RBC relaxation time *τ*_*µ*_ = *η*_*o*_*D*_*r*_*/µ*_*r*_ with membrane shear modulus *µ*_*r*_ and viscosity of the suspending medium *η*_*o*_. Another frequently used RBC relaxation time is based on the membrane’s bending rigidity *κ*_*r*_ and defined as 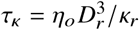. The cytosol viscosity *η*_*i*_ inside a RBC is about five times larger than the plasma viscosity (35), and the membrane’s shear viscosity is not considered here. The RBC relaxation time *τ*_*µ*_ is equal to 1.22 ms (compare to *τ*_*κ*_ = 0.86 s) for *η*_*o*_ = 9 × 10^−4^ Pa·s (a water-like PBS solution at 24 °C) and average properties of a healthy RBC, corresponding to *D*_*r*_ = 6.53 *µ*m, *µ*_*r*_ = 4.8 × 10^−6^ N/m, and *κ*_*r*_ = 70 *k*_*B*_*T* = 2.9 × 10^−19^ J. The average Young’s modulus of a healthy RBC is equal to *Y*_*r*_ = 18.9 × 10^−6^ N/m (37). Another important parameter, which characterizes the relative importance of RBC shear elasticity to bending rigidity, is the Föppl-von Kármán number 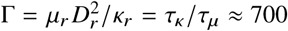.

**Table 1:**
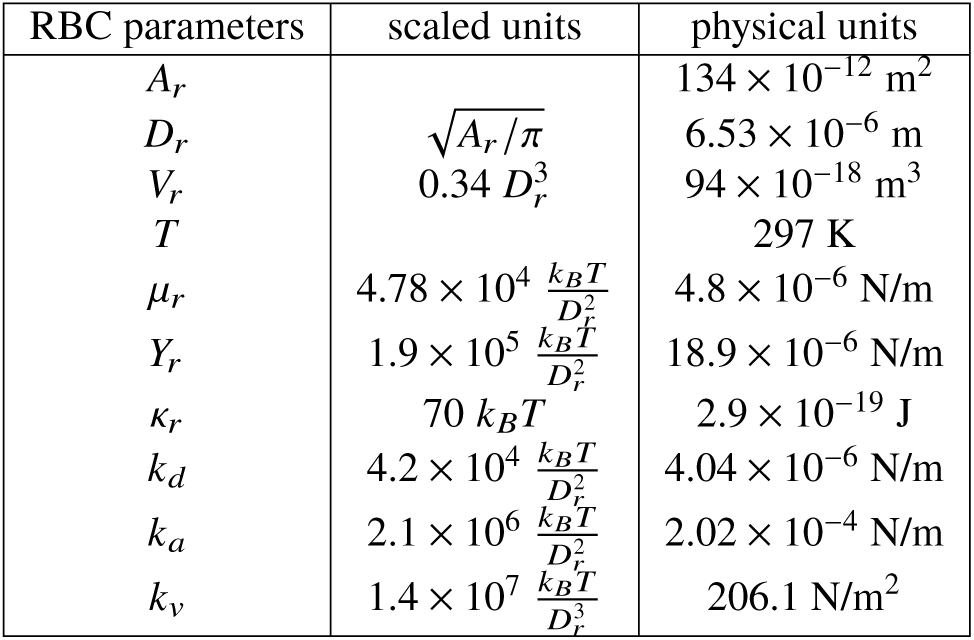
RBC parameters in units of the effective RBC diameter *D*_*r*_ and the thermal energy *k*_*B*_*T*, and the corresponding average values for a healthy RBC in physical units. *A*_*r*_ is the RBC membrane area, *V*_*r*_ is the RBC volume, *T* is the temperature, *µ*_*r*_ is the membrane shear modulus, *Y*_*r*_ is the Young’s modulus, *κ*_*r*_ is the membrane bending rigidity, and *k*_*d*_, *k*_*a*_, and *k*_*v*_ are the local area, global area, and volume constraint coefficients, respectively (37, 38). In all simulations, we have chosen *A*_*r*_ = 133.2 and *k*_*B*_*T* = 0.1, which implies that *D*_*r*_ = 6.51.

The membrane constitutes an impenetrable surface, which is modeled by bounce-back reflections of both cytosol and suspending-medium particles from inside and outside the membrane, respectively (37). Coupling between the RBC membrane and SDPD fluids is implemented through dissipative (friction) forces (37).

### Mesoscale simulations

To model fluid flow, we use the smoothed dissipative particle dynamics (SDPD) method (44) with angular momentum conservation (45) implemented by our group within the LAMMPS package (46). SDPD is a particle-based hydrodynamic technique employed to model the flow of both RBC cytosol and suspending medium. SDPD allows a direct input of fluid transport properties such as viscosity and of the equation of state for pressure, permitting a flexible control of fluid compressibility (47).

The SDPD fluid parameters are given in Table 2. Here, *p*_0_ and *b* are parameters of the equation of state *p* = *p*_0_(*ρ*/*ρ*_0_)^*α*^ −*b*, where *ρ* is the instantaneous particle density, *α* = 7, and *ρ*_0_ = *n* with *n* being fluid’s number density. Relatively large values of *p*_0_ and *α* provide a good approximation of fluid incompressibility, since the speed of sound *c* for this equation of state is *c*^2^ = *p*_0_*α*/*ρ*_0_.

**Table 2:** SDPD fluid parameters in simulation units. Mass and length for SDPD fluid are measured in units of the fluid particle mass *m* and the cutoff radius *r*_*c*_. *p*_0_ and *b* are parameters for the pressure equation, *η*_*o*_ is the dynamic viscosity of suspending fluid, and *n* is the number density. In all simulations, we have set *m* = 1, *r*_*c*_ = 1.0, and the thermal energy *k*_*B*_*T* = 0.1.

| Fluid parameters | Scaled units |
| --- | --- |
| $p_0$ | $1.5 \times 10^4 \frac{k_B T}{r_c^3}$ |
| $b$ | $1.49 \times 10^4 \frac{k_B T}{r_c^3}$ |
| $\eta_o$ | $80 - 365 \frac{\sqrt{mk_B T}}{r_c^2}$ |
| $n$ | $12 r_c^{-3}$ |

To span a wide range of flow rates, different values of suspending fluid viscosities 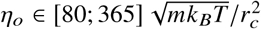 were employed in simulations (*m* is the fluid’s particle mass). Since the fluid viscosity modifies linearly the RBC relaxation time scale *τ*_*µ*_, large values of viscosity were used to model high flow rates in order to keep the Reynolds number 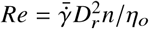 in simulations sufficiently low. Thus, in all simulations, *Re* = 0.2 was fixed, while *η*_*o*_ was computed for a desired flow rate 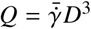, where *D* is the side length of a square channel.

### Simulation setup

A single RBC is suspended into a viscous fluid inside a square channel with a side length *D*, see simulation snapshots in Fig. 1. Periodic boundary conditions are assumed in the flow direction with a periodic length of *L* = 10*D*_*r*_ in all cases. The measurement of RBC shape is performed after the RBC passes this length numerous times, such that a final stationary state is achieved. The channel walls are modeled by frozen particles which assume the same structure as the fluid, while the wall thickness is equal to *r*_*c*_. To prevent wall penetration, fluid particles as well as vertices of a RBC are subject to bounce-back reflection at the fluid-solid interface. To ensure that no-slip boundary conditions are strictly satisfied at the walls, we also add a tangential adaptive shear force (48) which acts on the fluid particles in a near-wall layer of a thickness *r*_*c*_. The ratio of cytosol viscosity to that of suspending fluid is *λ* = *η*_*i*_ *η*_*o*_ = 5. The flow is driven by a force *f* applied to each fluid particle (both cytosol and external fluid), mimicking a pressure gradient Δ*P/L* = *fn*, where Δ*P* is the pressure drop. All simulations were performed on the supercomputer JURECA (49) at Forschungszentrum Jülich.

**Figure 1:**
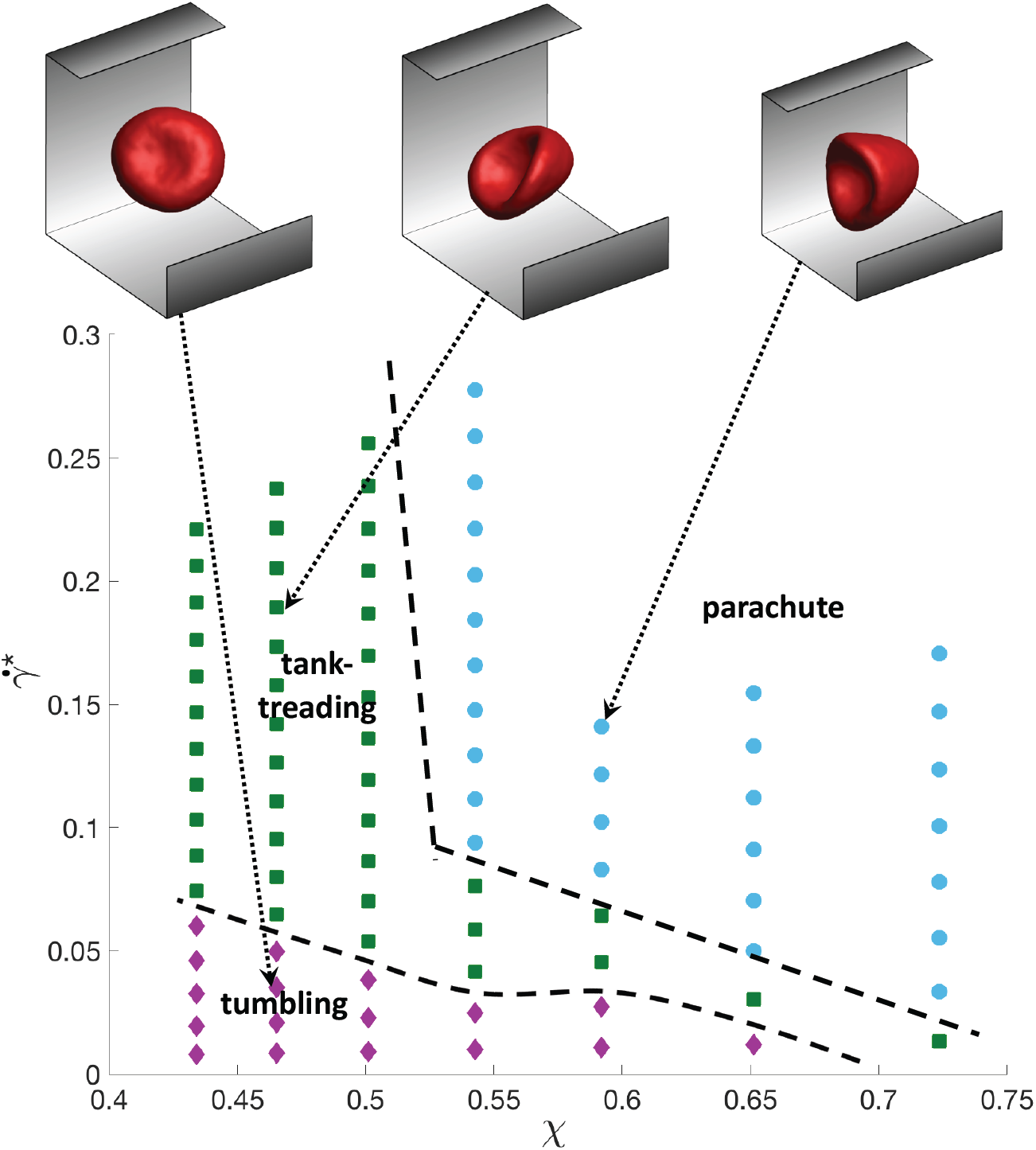
State diagram of RBC shapes and dynamics from simulations. Different shapes and dynamics of RBCs in square-channel flow for a wide range of confinements *χ* and non-dimensional shear rates 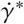. The RBC model represents average properties of a healthy RBC with *µ*_*r*_ = 4.8 × 10^−6^ N/m, *Y*_*r*_ = 18.9 × 10^−6^ N/m, *κ*_*r*_ = 2.9 × 10^−19^ J, and *λ* = *η*_*i*_ *η*_*o*_ = 5. The symbols correspond to separate simulations with different RBC states: parachute (cyan circle), tank-treading (green square), and tumbling (magenta diamond) cells. On the top, several representative shapes in flow are shown, see Movies S3-S5. Note that the state-boundary lines are drawn schematically to guide the eye.

### Blood samples

Blood was taken after informed consent from healthy donors by finger pricking with a lancet (Sarstedt Safety-Lancet Normal 21, Nürmbrecht). The study was approved by the Institutional Review Board of the University Medical Center Carl Gustav Carus at the Technische Universität Dresden (EK89032013, EK458102015). 0.5 – 1.5 *µ*l of whole blood was pipetted and suspended into 0.5 ml of PBS. The osmolality of the PBS was determined by an osmometer (The Fiske Micro-Osmometer Model 210, Norwood MA) at every measurement day and was found in the range of (313 ± 4) mOsm/kg (mean ± SD). Experimental measurements were performed at 24 °C within 30 minutes after blood drawing.

### Experimental setup

The experimental setup is similar to that previously described for real-time deformability cytometry (RT-DC) (50). Briefly, an Axiovert 200M (Zeiss, Oberkochen) inverted microscope was equipped with an EC Plan-NEOFLUAR 40x/0.75 objective (Zeiss, Oberkochen). To record the images, a MCI1362 CMOS camera (Mikrotron, Unterschleissheim) was connected to the microscope. The camera was controlled by the analysis software Shapeln (Zellmechanik Dresden, Dresden), which enables online cell tracking. The microscope was equipped with a custom-built illumination source using a high power blue LED (50). The LED was synchronized with the camera shutter for an exposure time of 2 *µ*s. Flow in the microfluidic-chip was initiated and controlled by a syringe pump (Cetoni neMESYS, Korbussen), which was connected to the chip with FEP-tubing. The flow rate of the cell suspension was controlled by a computer program provided by the manufacturer. For a series of experiments using a particular channel size, the range of flow rates was set such that the motion of cells with velocities between 0.1 mm/s and 4 mm/s could be observed.

Before starting experiments, the device was first filled with PBS from the sheath inlet at a flow rate of 1 *µ*l s. The chip was considered filled when the fluid reached the top of the sample inlet hole and the flow rate was then lowered to 0.1 *µ*l / s. This procedure prevents bubble formation when connecting the sample inlet tubing later. After connecting the sample tubing, the chip was filled with the RBC-suspension at 0.1 *µ*l / s from the sample tubing. When cells were observed flowing through the chip, the flow rate was lowered to 0.001 *µ*l / s for both sample and sheath flow and after 6 minutes waiting time for flow equilibration, cells were recorded. The online-tracking software developed for RT-DC (50) has been used to analyze and save images of RBCs. No continuous videos were recorded, in order to avoid numerous frames with an empty channel (no RBCs present). The software automatically stores and generates data about the position of a cell in the channel for each frame and other useful parameters such as size and aspect ratio.

### Experimental procedure

Diluted blood suspension (2 *µ*l whole blood in 0.5 ml PBS) was flown into the sample inlet, while the sheath inlet was fed with PBS. Starting from a sheath flow of 0.002 *µ*l / s, the flow rate was incrementally increased by 0.002 *µ*l / s up to a final total flow rate of 0.04 *µ*l / s. The sheath to sample flow ratio was kept 3 : 1 for all measurements.

Recording was done at a frame rate of 100 Hz for flow rates smaller than 0.006 *µ*l / s, 500 Hz for the range of 0.006 *µ*l / s −0.02 *µ*l / s, and 900 Hz for flow rates larger than 0.02 *µ*l /s. For each flow rate about 50 cells were recorded. For post analysis, the videos of the cells were separated in sections that contained the passage of only one cell, from which the mean velocity and shape classification were determined.

### Layout of the microfluidic chip

The syringe pump and the microfluidic chip layout typically used in RT-DC did not permit to achieve flow velocities small enough to observe non-deformed cells. Therefore, the channel system shown in Fig. 2*A* was designed to reduce the flow rate inside the measurement channel. Thus, the incoming total flow was split into three branches, where the main portion of the flow was diverted away into two outer branches and only a small fraction flows through the measurement channel. The widths of measurement channels and outer branches are given in Table S1. To increase the number of cells that reach the measurement channel, the cells were focused by a sheath flow before entering the region where the flow was split. The chip design from the inlets until the focusing channel was identical to that used for RT-DC (50). The cells that enter the measurement channel pass a serpentine region, which was introduced to prevent them from adhesion at the channel wall and to facilitate their entry position close to the channel center (see Movies S1 and S2). The measurement channel has a total length of 5.35 mm and measurements were performed approximately 4.5 mm after the entrance over a length of 435 *µ*m. This channel length was chosen to give the cells enough time to obtain a stable state. The devices were designed with the free software KLayout (51).

**Figure 2:**
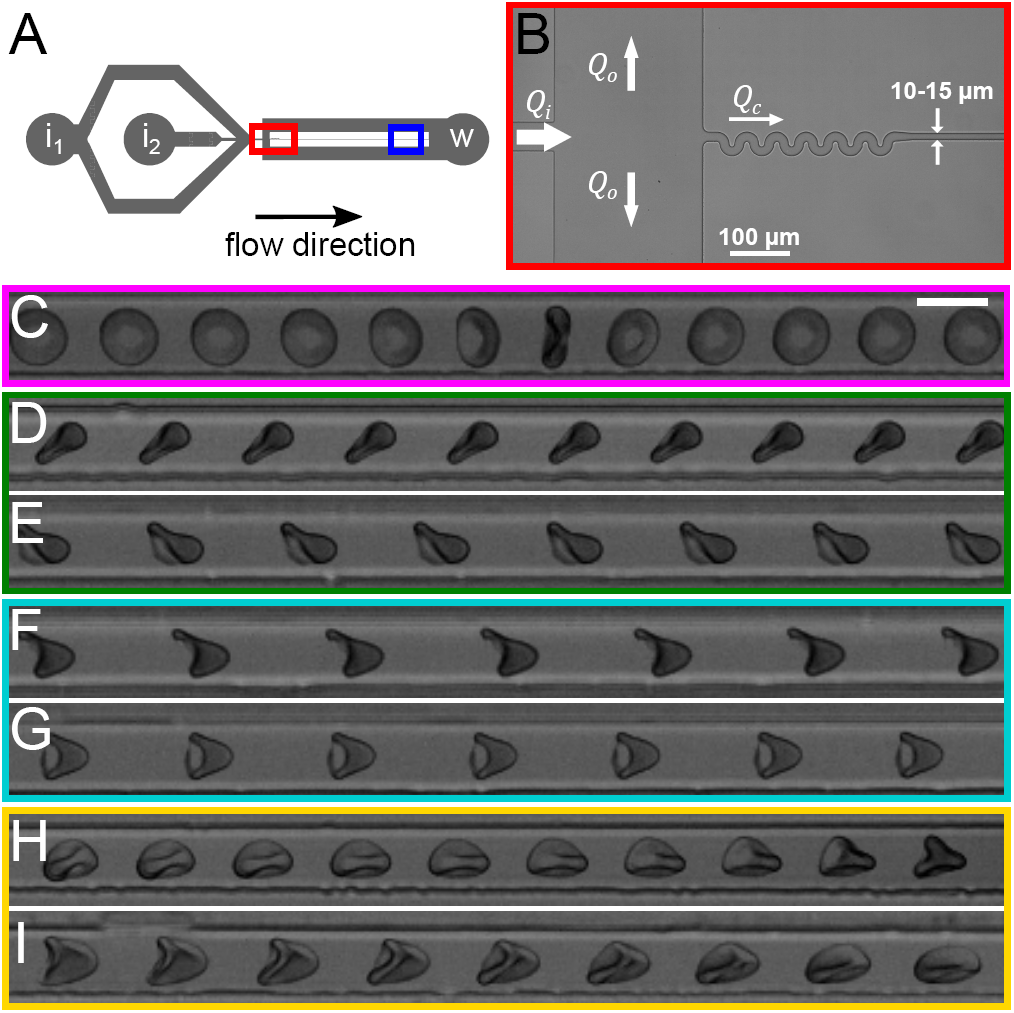
Microfluidic chip design and observed RBC states. (*A*) Complete layout of the channel system. Inlet i_1_ is for sheath flow, the sample inlet i_2_ supplies the blood cells and the w denotes the waste outlet. The region within the red rectangle is shown in more detail in *B*. The blue rectangle highlights the measurement region. (*B*) Section of the channel layout (marked by red rectangle in *A*) where the total flow *Q*_*i*_ is separated into channel flow *Q*_*c*_ and outer branch flow *Q*_*o*_. The arrow direction corresponds to the flow direction and the arrow width qualitatively represents the flow rate at a given point (not to scale). The serpentine at the channel entrance are necessary to center cells and avoid their attachment to the channel wall. (*C-I*) Examples of observed cell states in the measurement region. Images were created by minimum intensity projection from images of the same cell during channel passage. The scale bar in *C* is 10 *µ*m and applies to all images. The colored rectangles highlight the classes these cells are assigned to (see Figs. 1 and 3). (*C*) Tumbling cell for *χ* = 0.55 at 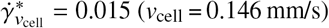, 75 ms between images. (*D*) Discocyte with a constant inclination angle for *χ* = 0.57 at 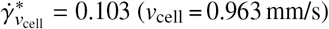, 15 ms between images. (*E*) Slipper for *χ* = 0.56 at 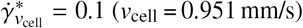, 20 ms between images. Cells as in *D* and *E* are classified as a tank-treading state. (*F*) Non-symmetrical parachute for *χ* = 0.56 at 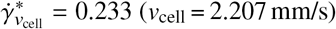, 10 ms between images. (*G*) Symmetrical parachute for *χ* = 0.56 at 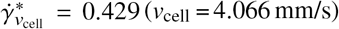, 5 ms between images. Cells as in *F* and *G* are classified as parachutes. (*H*) Tumbling polylobe for *χ* = 0.64 at 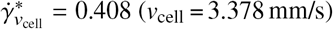, 5 ms between images. (*I*) Shape changing polylobe for *χ* = 0.59 at 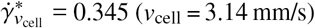, 5 ms between images. Cells as in *H* and *I* are classified as polylobes.

### Device fabrication

Microfluidic chips were made from PDMS for every experiment using standard soft lithography techniques and sealed with a glass cover slip after plasma surface activation. The aim was to achieve square cross sections with varying channel sizes. For this, it was necessary to produce a master on a silicon-wafer for every intended channel height. For the fabrication of the masters, first the silicon wafer was spin-coated with the photoresist AZ15nxT (The MicroChemicals GmbH, Ulm). A first spinning step of 10 s at 500 rpm (acceleration of 100 rpm/s) spreads the photoresist over the whole wafer. Then, the spinning frequency was increased in a second spinning step (acceleration of 300 rpm/s) to get a coating of the silicon with a thickness that equals the desired channel height on the finished chips. The frequency of the second spinning step for each wafer is given in Table S2.

After the wafers were coated, they were soft-baked for 5 minutes at 110 °C. In the next step, the coated wafer was exposed to UV-light (400 nm) with the photomask on top in contact mode, with an exposure power of 450 J/s. Then, the wafer was post-baked at 120 °C for 2 minutes to enhance cross-linking of the activated part of the photo-resist and the wafer. In a final step, the wafers were developed in a mixture of AZ400K developer (The MicroChemicals GmbH, Ulm) and water (1:2) for approximately 100 seconds. The developer dissolved the photo resist that was not activated. Finally, a hydrophobic coating (1H,1H,2H,2H-perfluorooctyl-trichlorosolane vapor (abcr, Germany)) was deposited onto the wafer under vacuum.

Even though microfluidic channels were designed to have a square cross-section, the fabrication of perfectly squared channels is difficult for such small systems. Therefore, channel sizes were characterized by a hydraulic diameter *D*_*h*_ = 4*A P*, where *A* is the channel’s cross-sectional area and *P* is the perimeter. The top width and the channel height were measured using a profilometer on the silicon wafers, which are used as masters in the production of the PDMS-microfluidic chips. The bottom width of the channel was measured with an inverted microscope by focusing on the bonding site of the PDMS and cover-glass. The width was then determined using a customized software. Because top and bottom widths of the channels did not always match, the cross-section was assumed as a symmetric trapezoid (see Fig. S1). The dimensions of all the channels used in the experiments can be found in Table S3.

### Data analysis

Each data point in state probability distributions is obtained as a weighted average probability of the measurements performed for the respective range of hydraulic diameters. For calculation of the average, every value was weighted by the number of cells *N*_*i*_ that were observed in the respective experiment *i*. Thus, experiments with a higher cell count have a higher weight w_*i*_ = *N*_*i*_/ Σ*j N*_*j*_. The weighted averages 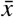 and corresponding standard deviations *σ* were calculated as 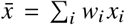 and 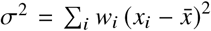. Here, *x*_*i*_ is the probability of a RBC state in the respective velocity range in experiment *i*. The error bars represent the weighted error for the set of data points.

## RESULTS

### Shape and dynamics diagram from simulations

To map RBC behavior for various conditions, a number of simulations with a periodic channel of square cross-section were performed for a wide range of channel sizes and flow rates. The side length *D* of a square channel determines RBC confinement, *χ* = *D*_*r*_*/D*. The flow rate was characterized by a non-dimensional shear rate as

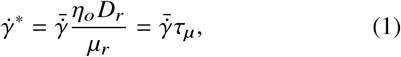

where 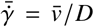 is the effective shear rate, 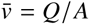 is the average flow velocity with cross-sectional area *A* = *D*^2^ and volumetric flow rate *Q*.

Figure 1 shows the simulated state diagram of RBC shapes and dynamics for a physiological viscosity contrast *λ* = 5. Three different states can be identified. At strong enough confinements and high flow rates, a parachute shape is observed, where the cell moves in the center of the channel and is deformed into a parachute-like shape by strong fluid forces. Here, we do not differentiate between symmetric and slightly asymmetric parachute shapes. At weak confinements, a RBC tumbles at low 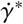 and attains a slipper shape at high 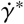. Both states are characterized by an off-center cell position. Slippers exhibit a nearly stationary orientation with tank-treading dynamics of the membrane. Hence, they will also be referred to as tank-treading cells. In the tumbling state, the cell exhibits typical tumbling motion without significant membrane tank-treading. It is important to note that for any fixed flow condition in simulations, a single state can generally be assigned with some exceptions very close to the state boundaries.

The diagram in Fig. 1 is qualitatively similar to the diagram for RBC behavior in tube flow at *λ* = 1 in Ref. (4). For a direct comparison, 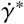 values in Fig. 1 should be multiplied by the Föppl-von Kármán number Γ ≈700, since the nondimensional shear rate in Ref. (4) has been defined as 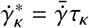. In case of *λ* = 5, the tumbling region is larger than for *λ* = 1 and expands toward larger 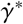 values. This is due to the fact that high viscosity contrasts suppress membrane tank-treading, thereby making the slipper state unfavorable. In fact, RBCs at *λ* = 5 in simple shear flow do not exhibit tank-treading motion at all (9, 36). However, under strong confinement, membrane tank-treading becomes possible, as we observe slippers in the diagram in Fig. 1, while under weak confinement no slipper shapes should be expected. An interesting result for *λ* = 5 in comparison to *λ* = 1 in Ref. (4) is that the parachute region shifts toward higher confinements. This is counter-intuitive in view of the argument that slippers should be suppressed at *λ* = 5. This result is consistent with 2D simulations for different viscosity contrasts (5, 34), which showed that slippers have their mass located closer to the centerline than parachutes, despite their position being slightly away from the channel center.

### RBC shapes from microfluidic experiments

For experiments, a microfluidic device depicted in Figs. 2*A* and 2*B* has been employed. At the beginning of the measurement channel, cells have to pass a serpentine region, so that cells which enter far off-center are forced to the channel center. The focusing effect of the serpentine region is due to the skewness of the flow profile in a bent section toward the inner wall with the larger curvature (52) and should not be confused with inertial focusing (53) found at high enough Reynolds numbers, because the characteristic Re in our experiments is smaller than 0.02. We observed that cells entering off-center primarily end up as slippers. This made it necessary to introduce the serpentine region to avoid that the way cells enter the channel influences the final state in the channel. Influence of the position at the channel entrance on the final state inside, especially the connection between starting off-center and ending up in a slipper shape, was recently described by Guckenberger et al. (33).

Cells were observed roughly 4.5 mm downstream of the channel entrance (blue region in Fig. 2*A*) to guarantee that cells have enough time to relax to a stable state, if there is one. The motion of each cell is characterized by average cell velocity *v*_cell_ in the measurement channel with a corresponding dimensionless shear rate

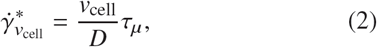

which is introduced in addition to 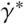, since the flow rate in the measurement channel cannot be directly accessed in experiments. Note that 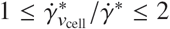. Typical cell states observed are shown in Figs. 2*C-I* and Movies S6-S12, and are divided into several classes. Figure 2*C* shows a nearly undeformed discocyte performing a flipping rotation in flow, which corresponds to the tumbling cells from the simulations. Discocyte-shaped RBCs also displayed a rolling motion or movement with a fixed inclination angle with respect to the flow axis, depicted in Fig. 2*D*. These states, which preserve the biconcave disk-like shape, mostly occur at low flow rates.

Increasing flow rate leads to deformation of the cells. The first mode of deformation is a bulging out of the front with a narrow tail at the rear of the cells, shown in Fig. 2*E*. Such cells usually have an off-center position and are classified as slippers. The states described in Figs. 2*D* and 2*E* are summarized as tank-treading RBCs, as they are similar to the tank-treading cells from the simulations, even though the membrane rotation cannot be directly observed experimentally. Further increase of the flow rate may lead to parachutes. Parachutes observed in the experiments were not always perfectly symmetrical. Therefore, a further distinction between asymmetrical (Fig. 2*F*) and symmetrical parachutes (Fig. 2*G*) was made. Still, both classes were summarized as parachutes for comparison with the simulations.

In wide enough channels at high flow rates, an additional state, a polylobe, was observed, which was not identified in simulations. Polylobe cells display multiple lobes and indentations, so that their general shape features do not fall into any of the other classes introduced before. Polylobes exhibited a nearly solid tumbling motion without significant shape changes (Fig. 2*H*) or strong shape changes which persisted during RBC passage (Fig. 2*I*). Polylobe shapes were observed at high shear rates in simple shear flow both experimentally and numerically (9, 36).

In experiments, multiple RBC states are always observed for a fixed flow condition for different cells. Therefore, experiments provide shape probability distributions as a function of 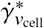, which are shown in Fig. 3*A* for channels with *χ* = 0.44 – 0.46 and *χ* = 0.64 – 0.67. Confinements *χ* were estimated by dividing the effective cell diameter *D*_*r*_ =6.53 *µ*m by the hydraulic diameter of the channel cross section. 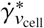 was binned in ranges of 0.025 from 0 to 0.425. Each data point is the weighted average probability of the measurements performed for the respective range of hydraulic diameters, see Materials and Methods for more details. The shape probability distributions in Fig. 3*A* clearly show that low confinement and 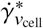 correspond primarily to tumbling RBCs. With increasing confinement, tank-treading becomes dominant at low 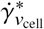 and parachutes take over with increasing cell velocity or 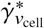. The probability for polylobes increases with 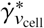 as well, but is only relevant for large channel sizes. The shape probability distributions for all investigated channel sizes are presented in Figs. S2 and S3.

**Figure 3:**
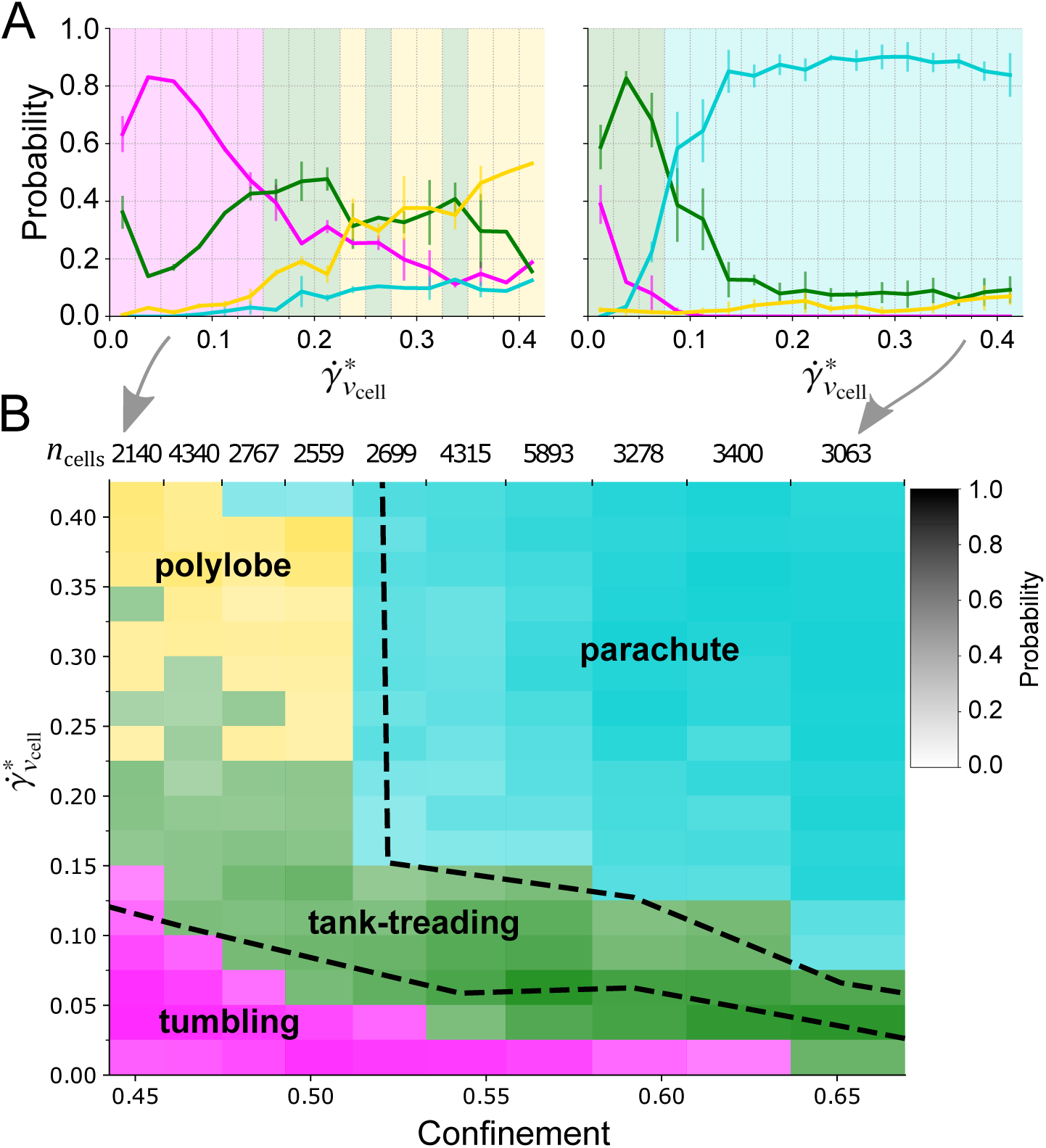
Experimental distribution of RBC states and comparison with simulation. (*A*) Velocity dependent probability distribution of shapes for the smallest (*χ* = 0.44 −0.46) and largest (*χ* = 0.64 −0.67) confinements from the experiments. The background shade highlights the shape with the highest probability in the corresponding velocity range. (*B*) State diagram of experimentally observed RBC states. The colors indicate the state with the highest probability within the corresponding confinement and velocity range. The saturation of the colors correlates with the probability to find the state. The dashed lines mark the state boundaries from the simulations; lower: tumbling to tank-treading, upper: tank-treading to parachute. The numbers above the graph show the number of cells that were classified for this range of confinements.

### Comparison between experiments and simulations

Experimental data were assembled into a state diagram shown in Fig. 3*B* as a function of 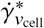 and the confinement, similarly to the simulated diagram in Fig. 1. The experimental diagram in Fig. 3*B* was constructed assuming the most probable shape. The saturation of the color indicates the probability for this state. The dashed lines represent the state boundaries from the simulations; the lower line corresponds to the limit between tumbling and tank-treading, while the upper line shows the boundary for tank-treading to parachute. The experimental diagram clearly shows that there are no sharp transitions between different states, as the probability to find the predominant state fades away the closer the state boundary is approached. A more detailed state diagram considering all subclasses introduced above can be found in Fig. S4.

Qualitatively, experimental and simulation diagrams in Figs. 3*B* and 1 are very similar. For small *χ* and 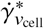 values, tumbling is the most likely state. With increasing 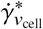 and *χ*, tank-treading becomes more apparent, and for large *χ* and 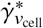 values, parachutes dominate.

However, there are several noticeable differences. For instance, the band, where tank-treading occurs, is narrower for the simulations in comparison to the experimental data. The differences in the state boundaries are likely due to different channel geometries and the variability in cell properties (e.g. size, bending rigidity, shear elasticity). Microfluidic channels in experiments did not have a perfect square geometry employed in simulations due to fabrication limitations, see Fig. S1 and Table S3. Even though the deviation from a perfect square geometry may not influence the distribution of shapes, it can affect the orientation of RBC shapes in the channel and therefore, their perception during experimental observations, as pointed out in Ref. (54). Furthermore, real RBCs are clearly not identical to each other and possess an appreciable variability in their size and mechanical properties, which will be discussed later. Another difference between the two diagrams is the existence of a polylobe region in experiments, which is absent in simulations. A probable reason is a difference in viscosity contrast *λ*. In simulations, a physiological viscosity contrast of *λ* = 5 at 37 °C was targeted, while experiments likely have a larger *λ* value. Note that the viscosity of blood plasma is larger than the viscosity of PBS solution used as a fluid medium. Furthermore, the viscosity of the RBC cytosol is sensitive to temperature changes and significantly increases at room temperature in comparison to the physiological temperature (55). Additionally, the presence of membrane viscosity, which was not modeled in simulations, would further increase the viscous resistance for cell deformation (56, 57). A high viscosity contrast is known to prohibit tank-treading motion of vesicles (58, 59) and RBCs (9, 36) in pure shear flow. Similarly, RBC tank-treading in a channel may be hindered by a high enough viscosity contrast, resulting in a polylobe.

Another difference between simulations and experiments is that the cells in the experiments may not always enter the channel perfectly centered, while a central cell positioning in the channel is the initial condition in simulations. The serpentine region in the experimental setup aids to bring RBCs close to the channel center, but of course it does not guarantee it. Nevertheless, the observations of RBC shapes are made about 4 mm downstream of the serpentine region, which is likely enough to achieve a stable RBC state, as channel sizes are comparable to the RBC diameter. Furthermore, the good agreement between simulations and experiments indicates that the entrance position of RBCs does not significantly affect observed RBC shapes.

Finally, it is worthwhile to mention that a small fraction of cells showed state changes within the region of observation. For instance, some RBCs entering the observation region as slippers or parachutes left it as a tumbling polylobe (see Movie S13). Similarly, RBCs entering as polylobes obtained a stable position in the channel and switched to a slipper or parachute within the observation window (see Movie S14). These examples suggest that there might be ranges of flow conditions and RBC mechanical properties, where two different states are quasi-stable and can interchange. Another possibility is that small perturbations in the flow due to geometrical irregularities of the channel trigger a state change. Interestingly, such state changes are mainly observed for channel sizes and flow rates close to the transitions between different shape and dynamics states.

### Approximation of RBC variability

Despite the good agreement between simulation and experimental diagrams in Fig. 3, the main dissimilarity is the existence of multiple RBC states for fixed flow conditions in experiments, while simulations generally yield a single state for fixed conditions. We attribute this result to the variability in mechanical properties of RBCs. Here, different RBC sizes (roughly within 6.5 – 9 *µ*m), RBC shear elasticities (*µ* ∈ [2, 10] *µ*N /m), membrane bending rigidities as well as cytosol and membrane viscosities may affect the observed state.

To investigate this proposition, we construct an empirical distribution for the variability of RBC properties and use it to estimate the frequency of appearance of different RBC states for fixed flow conditions. Since we have not investigated numerically the dependence of RBC behavior on bending rigidity and the cytosol and membrane viscosities, the effect of their variability cannot be studied here. Furthermore, we focus only on the variability in membrane shear elasticity and omit any variations in RBC size, so that a one-dimensional distribution is considered. Figure 4*A* shows a sample distribution of RBC shear elasticity. To construct this distribution, several assumptions are made. We consider blood sample to be a mixture of RBCs of various ages within the total lifespan of 120 days. About 2 × 10^11^ new RBCs are produced per day, whose shear modulus is assumed to have a Gaussian distribution with a mean *µ*_young_ = 5.5 *µ*N /m (60, 61) and standard deviation *σ* = 2 *µ*N /m (*σ* is an adjustable parameter). During aging, RBCs become stifer. The stifening process is assumed to have a linear temporal dependence of the shear modulus, which is represented by a linear shift of the Gaussian distribution for new RBCs toward larger shear moduli, such that after 120 days the mean of the distribution becomes *µ*_old_ = 6 *µ*N /m (60, 61). Then, the stiffness distribution for a blood sample in Fig. 4*A* corresponds to a superposition of Gaussian distributions of RBCs of different ages.

**Figure 4:**
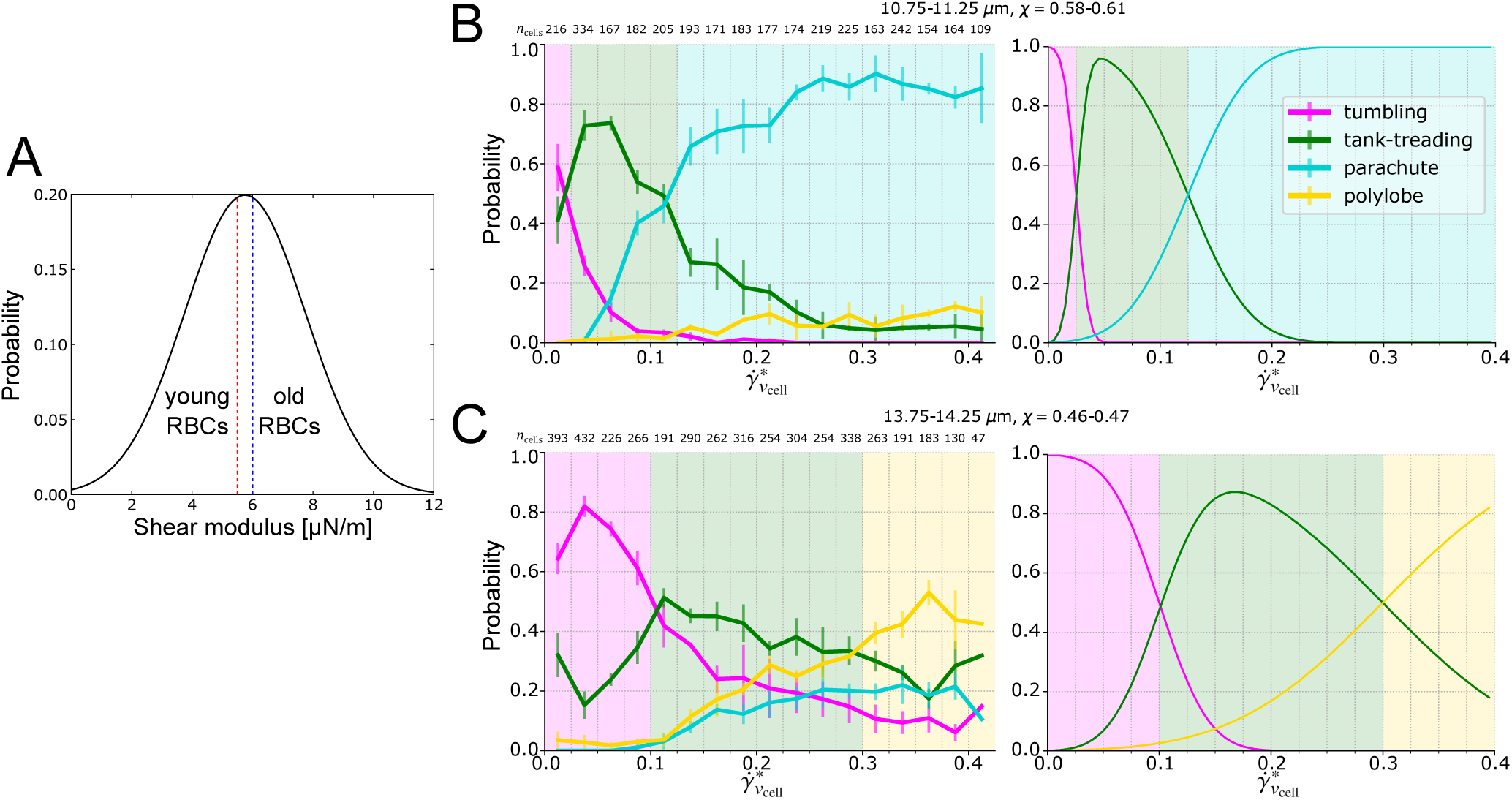
Characterization of RBC variability. (*A*) Distribution of the stiffness (shear modulus) of RBCs over their lifetime. (*B,C*) Comparison of the probabilities from experiment and approximation of RBC variability for different RBC states using the assumed distribution for (*B*) *χ* = 0.58 −0.61 and (*C*) *χ* = 0.46 −0.47. Left: experiment, right: theory. The numbers above the graph show the number of cells that were classified for this range of velocities.

Figure 3*B* shows the comparison of simulation and experimental data for RBC states as a function of 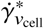, which is implicitly a function of *v*_cell_ /*µ*_*r*_. Under the assumption that 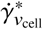 is the main parameter governing RBC behavior for a fixed *χ*, we obtain an equivalence between cell velocity (or flow rate) and RBC stiffness, i.e. slow flows and soft cells are equivalent to fast flows and stiff cells. Therefore, without loss of generality, we can assume that changes in 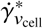 in Fig. 3*B* also represent corresponding changes in the cell stiffness characterized by *µ*_*r*_. As a result, for any selected point in the state diagram in Fig. 3*B*, we can superimpose the shear-modulus distribution in Fig. 4*A* onto the diagram by placing the peak of the distribution at this point (see schematic in Fig. S5). Then, the state boundaries in Fig. 3*B* typically divide the shear-modulus distribution into three regimes: (i) tumbling (tb) 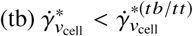, (ii) tank-treading (tt) 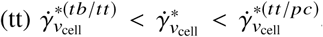, and (iii) parachute 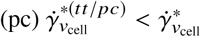; the corresponding integrated parts of the probability distribution provide the probability for observing tumbling, tank-treading, and parachute states, respectively. Note that these arguments apply within regime of linear elasticity theory.

Predicted state distributions in comparison to those measured experimentally are shown in Figs. 4*B* and 4*C* for two confinements as a function of 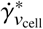. The qualitative agreement is very good, indicating that the variability in mechanical properties is indeed the primary reason for the observation of multiple RBC states in experiments for fixed flow conditions. The behavior of probability curves in Figs. 4*B* and 4*C* is directly associated with the width *σ* of shear-modulus distribution in Fig. 4*A*. Thus, a narrower distribution in shear modulus would result in a sharper crossover from one state to the other, while a wider distribution in *µ* can significantly broaden the crossover.

Clearly, the main limitation of the predicted distributions is that they are based only on the variation in membrane shear modulus and do not consider other possible differences between RBCs. In general, variations in RBC properties should be represented by a multidimensional distribution, which can be superimposed onto a multidimensional state diagram with axes representing different RBC properties. However, the good agreement between the shape distributions based on the proposed variation in shear modulus and those directly-measured experimentally indicates that the shear modulus is one of the main parameters governing RBC shapes in microchannel flow.

## DISCUSSION

Even though a number of previous experimental studies (27–31, 33) have demonstrated the existence of different RBC states (e.g. parachutes, slippers), our experimental investigation is the first where different RBC states are systematically mapped into a diagram as a function of channel size and flow rate. This was possible due to a large number of experimental observations, as about 35 000 cells were processed. In addition, present experiments directly demonstrate the existence of new RBC states in Poiseuille flow, such as tumbling and polylobe, not observed previously in microfluidic channels. Tumbling discocytes have been predicted in simulations (4) and now confirmed experimentally, even though such dynamics is not very intuitive, as RBCs tumble in Poiseuille flow in channels with sizes comparable to the RBC diameter. Simulations show that a tumbling cell assumes an off-center position, which results in asymmetrical shear stresses rotating the cell. Furthermore, RBC tumbling occurs at relatively low flow rates, at which fluid stresses are too weak to deform the RBC into a parachute or slipper. In the experiments, the cells at low flow rates may take up an off-center position due to sedimentation within the channel. At high flow rates, experimental observations demonstrate the existence of polylobe shapes, observed recently in pure shear flow (9, 36), also in channels wider than about 13 *µ*m.

In addition to the RBC state diagram, our microfluidic experiments show that there is no single state for fixed flow conditions, but rather a distribution of different cell states for each flow rate. In contrast, simulations yield primarily a single RBC state for a fixed confinement and flow rate. This result can be clearly attributed to the variability in RBC properties, which is inherently present in RBC samples. A simplified approximation for the variability in RBC shear elasticity demonstrates a good correspondence between experimentally observed distributions and those obtained by the superposition of the proposed variability distribution onto the state diagram. This reveals the complexity in real blood samples, especially if quantitative predictions of cells’ mechanical properties are aimed for. In addition to the variation in shear elasticity, variability in cell size, membrane bending rigidity, and cytosol and membrane viscosities are likely to have an effect on measured distributions of RBC states. To take these parameters into consideration, a detailed understanding of the effect of these properties on the RBC state diagram is required.

The good agreement between the simulated and experimental diagrams in Fig. 3 allows us to make a step toward the quantification of experimental observations. Reliable simulation data can be used to determine mechanical properties of flowing RBCs. As an example, for a fixed flow rate, the fraction of RBCs whose shear modulus is smaller/larger than a particular value can be determined by a state boundary between different RBC states. Thus, by carefully selecting flow rate and cell confinement (or channel size), specific values of RBC shear elasticity can be targeted and softer/stiffer cells can be differentiated through the observation of their shapes and dynamics in channel flow. This can be useful in blood analysis, since RBC deformability might be altered in many blood related diseases and disorders, such as malaria, sickle-cell anemia, diabetes, etc. (13, 14, 21).

Data presented in Fig. 3 also exhibit appreciable differences between simulations and experiments. The differences in state boundaries likely come from the different channel geometries (a square channel in simulations and a trapezoidal channel in experiments) and the variability in cell properties. Another difference is that tumbling RBCs and slippers generally display more deformation in simulations than in experiments. Also, experiments demonstrated the existence of polylobe shapes, which have not been observed in simulations. These discrepancies indicate that further improvements/ additions might be needed for the simulation model to fully capture experimental observations. A recent simulation study (33) suggests that RBCs might have bistable states in a channel under certain conditions determined by flow velocity and the cells’ initial position in the channel. In our experiments, the initial position and orientation of RBCs as they enter the channel have some variability and cannot be precisely controlled. Considering the results from Ref. (33), it can be expected that the probability to find tank-treading cells in the experiment would decrease if all cells would start at a perfectly centered position. Another limitation of experiments is that the observations are made within a window of about 435 *µ*m in length, which cells pass relatively fast. The performed simulations indicate that the full period of cell dynamics (e.g. when the membrane of a tank-treading cell performed half of its full rotation) is often about several times longer than the observation times in experiments. This means that only a portion of cell dynamics is directly monitored for every single cell, which prevents the complete elimination of the existence of dynamic deformations observed for tumbling RBCs and slippers in simulations.

## CONCLUSIONS

In summary, our combined simulation and experimental investigation of RBC shapes and dynamics in microchannels provides a consistent RBC state diagram and illustrates the complexity of RBC behavior in microflow. The simulation results agree well with experimental observations, allow the characterization of RBC variability in shear elasticity, and permit us to make a significant step toward quantitative measurements of RBC mechanical properties. These results are expected to encourage further numerical and microfluidic investigations of RBC behavior in microchannels or microvessels in order to better understand how various properties of RBCs affect their behavior in microflow and to advance toward reliable and fast detection of changes in RBC deformability relevant in many blood diseases and disorders.

## AUTHOR CONTRIBUTIONS

F.R. performed experiments. J.M. and D.A.F. performed simulations. A.A.N. fabricated silicon masters for microfluidic chip production. F.R., J.M., and D.A.F. analyzed data. G.G., J.G., and D.A.F. designed research. All authors discussed the results and wrote the paper.

## ACKNOWLEDGMENTS

D.A.F. and J.G. acknowledge funding by the Alexander von Humboldt Foundation. We acknowledge funding from the FP7 ITN “LAPASO” to J.G., G.G. and D.A.F. We also gratefully acknowledge the computing time granted through JARA-HPC on the supercomputer JURECA at Forschungszentrum Jülich. We thank Salvatore Girardo and the Microstructure Facility of the CMCB (in part funded by the State of Saxony and the European Fund for Regional Development -EFRE) for assistance with the production of Si-wafers, and Oliver Otto, Philipp Rosendahl, Martin Kräter, and Maik Herbig for their input regarding this project. We gratefully thank Nicole Toepfner who wrote the ethics proposals for this project.

## Notes

#### Summary of Updates

Final script accepted by journal Author information updated

## REFERENCES

1. Popel, A. S., and P. C. Johnson, 2005. Microcirculation and hemorheology. Annu. Rev. Fluid Mech. 37:43–69.

2. Lipowsky, H. H., 2005. Microvascular rheology and hemodynamics. Microcirculation 12:5–15.

3. Noguchi, H., and G. Gompper, 2005. Shape transitions of fluid vesicles and red blood cells in capillary flows. Proc. Natl. Acad. Sci. USA 102:14159–14164.

4. Fedosov, D. A., M. Peltomäki, and G. Gompper, 2014. Deformation and dynamics of red blood cells in flow through cylindrical microchannels. Soft Matter 10:4258–4267.

5. Kaoui, B., N. Tahiri, T. Biben, H. Ez-Zahraouy, A. Benyoussef, G. Biros, and C. Misbah, 2011. Complexity of vesicle microcirculation. Phys. Rev. E 84:041906.

6. Wells Jr, R. E., and E. W. Merrill, 1961. Shear rate dependence of the viscosity of whole bllod and plasma. Science 133:763–764.

7. Skalak, R., S. R. Keller, and T. W. Secomb, 1981. Mechanics of blood flow. J. Biomech. Eng. 103:102–115.

8. Fedosov, D. A., W. Pan, B. Caswell, G. Gompper, and G. E. Karniadakis, 2011. Predicting human blood viscosity in silico. Proc. Natl. Acad. Sci. USA 108:11772–11777.

9. Lanotte, L., J. Mauer, S. Mendez, D. A. Fedosov, J.-M. Fromental, V. Claveria, F. Nicoud, G. Gompper, and M. Abkarian, 2016. Red cells’ dynamic morphologies govern blood shear thinning under microcirculatory flow conditions. Proc. Natl. Acad. Sci. USA 113:13289–13294.

10. Kon, K., N. Maeda, and T. Shiga, 1983. The influence of deformation of transformed erythrocytes during flow on the rate of oxygen release. J. Physiol. 339:573–584.

11. Sprague, R. S., M. L. Ellsworth, A. H. Stephenson, M. E. Kleinhenz, and A. J. Lonigro, 1998. Deformation-induced ATP release from red blood cells requires CFTR activity. Am. J. Physiol. 275:H1726–H1732.

12. Forsyth, A. M., J. Wan, P. D. Owrutsky, M. Abkarian, and H. A. Stone, 2011. Multiscale approach to link red blood cell dynamics, shear viscosity, and ATP release. Proc. Natl. Acad. Sci. USA 108:10986–10991.

13. Diez-Silva, M., M. Dao, J. Han, C.-T. Lim, and S. Suresh, 2010. Shape and biomechanical characteristics of human red blood cells in health and disease. MRS Bulletin 35:382–388.

14. Fedosov, D. A., M. Dao, G. E. Karniadakis, and S. Suresh, 2014. Computational biorheology of human blood flow in health and disease. Ann. Biomed. Eng. 42:368–387.

15. Kaul, D. K., M. E. Fabry, P. Windisch, S. Baez, and L. Nagel, 1983. Erythrocytes in sickle cell anemia are heterogeneous in their rheological and hemodynamic characteristics. J. Clin. Invest. 72:22–31.

16. Barabino, G. A., M. O. Platt, and D. K. Kaul, 2010. Sickle cell biomechanics. Annu. Rev. Biomed. Eng. 12:345–367.

17. Tsukada, K., E. Sekizuka, C. Oshio, and H. Minamitani, 2001. Direct measurement of erythrocyte deformability in diabetes mellitus with a transparent microchannel capillary model and high-speed video camera system. Microvasc. Res. 61:231–239.

18. Brown, C. D., H. S. Ghali, Z. Zhao, L. L. Thomas, and E. A. Friedman, 2005. Association of reduced red blood cell deformability and diabetic nephropathy. Kidney Int. 67:295–300.

19. Cranston, H. A., C. W. Boylan, G. L. Carroll, S. P. Sutera, J. R. Williamson, I. Y. Gluzman, and D. J. Krogstad, 1984. Plasmodium falciparum maturation abolishes physiologic red cell deformability. Science 223:400–403.

20. Shelby, J. P., J. White, K. Ganesan, P. K. Rathod, and D. T. Chiu, 2003. A microfluidic model for single-cell capillary obstruction by Plasmodium falciparum-infected erythrocytes. Proc. Natl. Acad. Sci. USA 100:14618–14622.

21. Koch, M., K. E. Wright, O. Otto, M. Herbig, N. D. Salinas, N. H. Tolia, T. J. Satchwell, J. Guck, N. J. Brooks, and J. Baum, 2017. Plasmodium falciparum erythrocyte-binding antigen 175 triggers a biophysical change in the red blood cell that facilitates invasion. Proc. Natl. Acad. Sci. USA 114:4225–4230.

22. Davis, J. A., D. W. Inglis, K. J. Morton, D. A. Lawrence, R. Huang, S. Y. Chou, J. C. Sturm, and R. H. Austin, 2006. Deterministic hydrodynamics: taking blood apart. Proc. Nat. Acad. Sci. USA 103:14779–14784.

23. Toner, M., and D. Irimia, 2005. Blood-on-a-chip. Annu. Rev. Biomed. Eng. 7:77–103.

24. Henry, E., S. H. Holm, Z. Zhang, J. P. Beech, J. O. Tegenfeldt, D. A. Fedosov, and G. Gompper, 2016. Sorting cells by their dynamical properties. Sci. Rep. 6:34375.

25. Toepfner, N., C. Herold, O. Otto, P. Rosendahl, A. Jacobi, Kräter, J. Stächele, L. Menschner, M. Herbig, L. Ciuffreda, L. Ranford-Cartwright, M. Grzybek, U. Coskun, E. Reithuber G. Garriss, P. Mellroth, B. Henriques-Normark, N. Tregay, M. Suttorp, M. Bornhäuser, E. R. Chilvers, R. Berner, and J. Guck, 2018. Detection of human disease conditions by single-cell morpho-rheological phenotyping of blood. eLife 7:e29213.

26. Dodgson, S. E., 2018. There will be blood tests. Cell 173:1–3.

27. Skalak, R., and P. I. Branemark, 1969. Deformation of red blood cells in capillaries. Science 164:717–719.

28. Gaehtgens, P., C. Dührssen, and K. H. Albrecht, 1980. Motion, deformation, and interaction of blood cells and plasma during flow through narrow capillary tubes. Blood Cells 6:799–812.

29. Bagge, U., P.-I. Branemark, R. Karlsson, and R. Skalak, 1980. Three-dimensional observations of red blood cell deformation in capillaries. Blood Cells 6:231–237.

30. Abkarian, M., M. Faivre, R. Horton, K. Smistrup, C. A. Best-Popescu, and H. A. Stone, 2008. Cellular-scale hydrodynamics. Biomed. Mater. 3:034011.

31. Tomaiuolo, G., M. Simeone, V. Martinelli, B. Rotoli, and S. Guido, 2009. Red blood cell deformation in microconfined flow. Soft Matter 5:3736–3740.

32. McWhirter, J. L., H. Noguchi, and G. Gompper, 2009. Flow-induced clustering and alignment of vesicles and red blood cells in microcapillaries. Proc. Natl. Acad. Sci. USA 106:6039–6043.

33. Guckenberger, A., A. Kihm, T. John, C. Wagner, and Gekle, 2018. Numerical-experimental observation of shape bistability of red blood cells flowing in a microchannel. Soft Matter 14:2032–2043.

34. Tahiri, N., T. Biben, H. Ez-Zahraouy, A. Benyoussef, and C. Misbah, 2013. On the problem of slipper shapes of red blood cells in the microvasculature. Microvasc. Res. 85:40–45.

35. Wells, R., and H. Schmid-Schönbein, 1969. Red cell deformation and fluidity of concentrated cell suspensions. J. Appl. Physiol 27:213–217.

36. Mauer, J., S. Mendez, L. Lanotte, F. Nicoud, M. Abkarian, G. Gompper, and D. A. Fedosov, 2018. Flow-induced transitions of red blood cell shapes under shear. Phys. Rev. Lett. 121:118103.

37. Fedosov, D. A., B. Caswell, and G. E. Karniadakis, 2010. A multiscale red blood cell model with accurate mechanics, rheology, and dynamics. Biophys. J. 98:2215–2225.

38. Fedosov, D. A., B. Caswell, and G. E. Karniadakis, 2010. Systematic coarse-graining of spectrin-level red blood cell models. Comput. Meth. Appl. Mech. Eng. 199:1937–1948.

39. Peng, Z., A. Mashayekh, and Q. Zhu, 2014. Erythrocyte responses in low-shear-rate flows: effects of non-biconcave stress-free state in the cytoskeleton. J. Fluid Mech. 742:96–118.

40. Cordasco, D., A. Yazdani, and P. Bagchi, 2014. Comparison of erythrocyte dynamics in shear flow under different stress-free configurations. Phys. Fluids 26:041902.

41. Abkarian, M., M. Faivre, and A. Viallat, 2007. Swinging of red blood cells under shear flow. Phys. Rev. Lett. 98:188302.

42. Dupire, J., M. Socol, and A. Viallat, 2012. Full dynamics of a red blood cell in shear flow. Proc. Natl. Acad. Sci. USA 109:20808–20813.

43. Sinha, K., and M. D. Graham, 2015. Dynamics of a single red blood cell in simple shear flow. Phys. Rev. E 92:042710.

44. Español, P., and M. Revenga, 2003. Smoothed dissipative particle dynamics. Phys. Rev. E 67:026705.

45. Müller, K., D. A. Fedosov, and G. Gompper, 2015. Smoothed dissipative particle dynamics with angular momentum conservation. J. Comp. Phys. 281:301–315.

46. Plimpton, S., 1995. Fast parallel algorithms for shortrange molecular dynamics. J. Chem. Phys. 117:1–19.

47. Alizadehrad, D., and D. A. Fedosov, 2018. Static and dynamic properties of smoothed dissipative particle dynamics. J. Comp. Phys. 356:303–318.

48. Fedosov, D. A., and G. E. Karniadakis, 2009. Triple-decker: Interfacing atomistic-mesoscopic-continuum flow regimes. J. Comp. Phys. 228:1157–1171.

49. Jülich Supercomputing Centre, 2018. JURECA: Modular supercomputer at Jülich Supercomputing Centre. J. Large-Scale Res. Facil. 4:A132.

50. Otto, O., P. Rosendahl, A. Mietke, S. Golfier, C. Herold Klaue, S. Girardo, S. Pagliara, A. Ekpenyong, A. Jacobi, M. Wobus, N. Töpfner, U. F. Keyser, J. Mansfeld, Fischer-Friedrich, and J. Guck, 2015. Real-time deformability cytometry: on-the-fly cell mechanical phenotyping. Nat. Methods 12:199–202.

51. Otto, O. 2006. KLayout - High Performance Layout Viewer And Editor. http://www.klayout.de/index.php.

52. Ye, T., N. Phan-Thien, C. T. Lim, and Y. Li, 2017. Red blood cell motion and deformation in a curved microvessel. J. Biomech. 65:12–22.

53. Di Carlo, D., D. Irimia, R. G. Tompkins, and M. Toner, 2007. Continuous inertial focusing, ordering, and separation of particles in microchannels. Proc. Natl. Acad. Sci. USA 104:18892–18897.

54. Coupier, G., A. Farutin, C. Minetti, T. Podgorski, and C. Misbah, 2012. Shape diagram of vesicles in Poiseuille flow. Phys. Rev. Lett. 108:178106.

55. Kelemen, C., S. Chien, and G. M. Artmann, 2001. Temperature transition of human hemoglobin at body temperature: effects of calcium. Biophys. J. 80:2622–2630.

56. Tran-Son-Tay, R., S. P. Sutera, and P. R. Rao, 1984. Determination of red blood cell membrane viscosity from rheoscopic observations of tank-treading motion. Biophys. J. 46:65–72.

57. Noguchi, H., and G. Gompper, 2004. Fluid vesicles with viscous membranes in shear flow. Phys. Rev. Lett. 93:258102.

58. Kantsler, V., and V. Steinberg, 2006. Transition to tumbling and two regimes of tumbling motion of a vesicle in shear flow. Phys. Rev. Lett. 96:036001.

59. Mader, M.-A., V. Vitkova, M. Abkarian, A. Viallat, and T. Podgorski, 2006. Dynamics of viscous vesicles in shear flow. Eur. Phys. J. E 19:389–397.

60. Nash, G. B., and S. J. Wyard, 1981. Erythrocyte membrane elasticity during in vivo ageing. Biochimica et Biophysica Acta 643:269–275.

61. Nash, G. B., and H. J. Meiselman, 1983. Red cell and ghost viscoelasticity. Effects of hemoglobin concentration and in vivo aging. Biophys. J. 43:63–73.

