## Supplementary material for "High-throughput microfluidic characterization of erythrocyte shapes and mechanical variability": SI text, tables, and figures

### SUPPLEMENTARY FIGURES

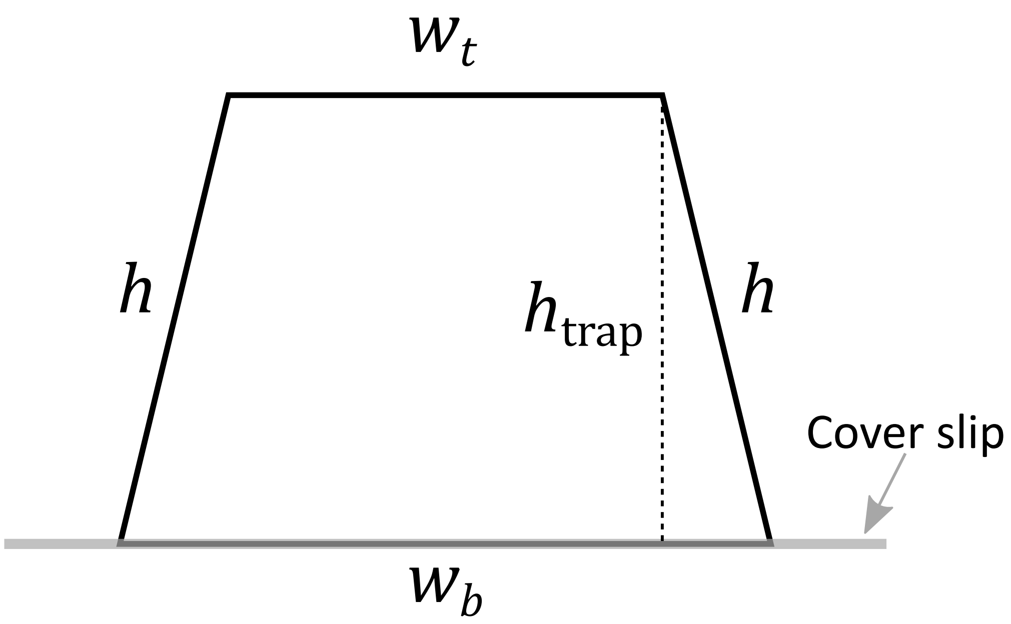


**FIGURE S1.** Schematic of the channel cross-section assumed as trapezoid. $w_t$ and $h$ are measured on the Si-master. It is assumed that the side walls are bent when bonding PDMS-chip to the cover slip. $w_b$ is measured on the chip (see Materials and Methods). The hydraulic diameter $D_{\text{hyd}} = 4A/P$ ($A$ - area of cross section, $P$ - trapezoid perimeter) is calculated with the following formula $D_{\text{hyd}} = \frac{2\cdot(w_t+w_b)\cdot h_{\text{trap}}}{w_t+w_b+2h}$.

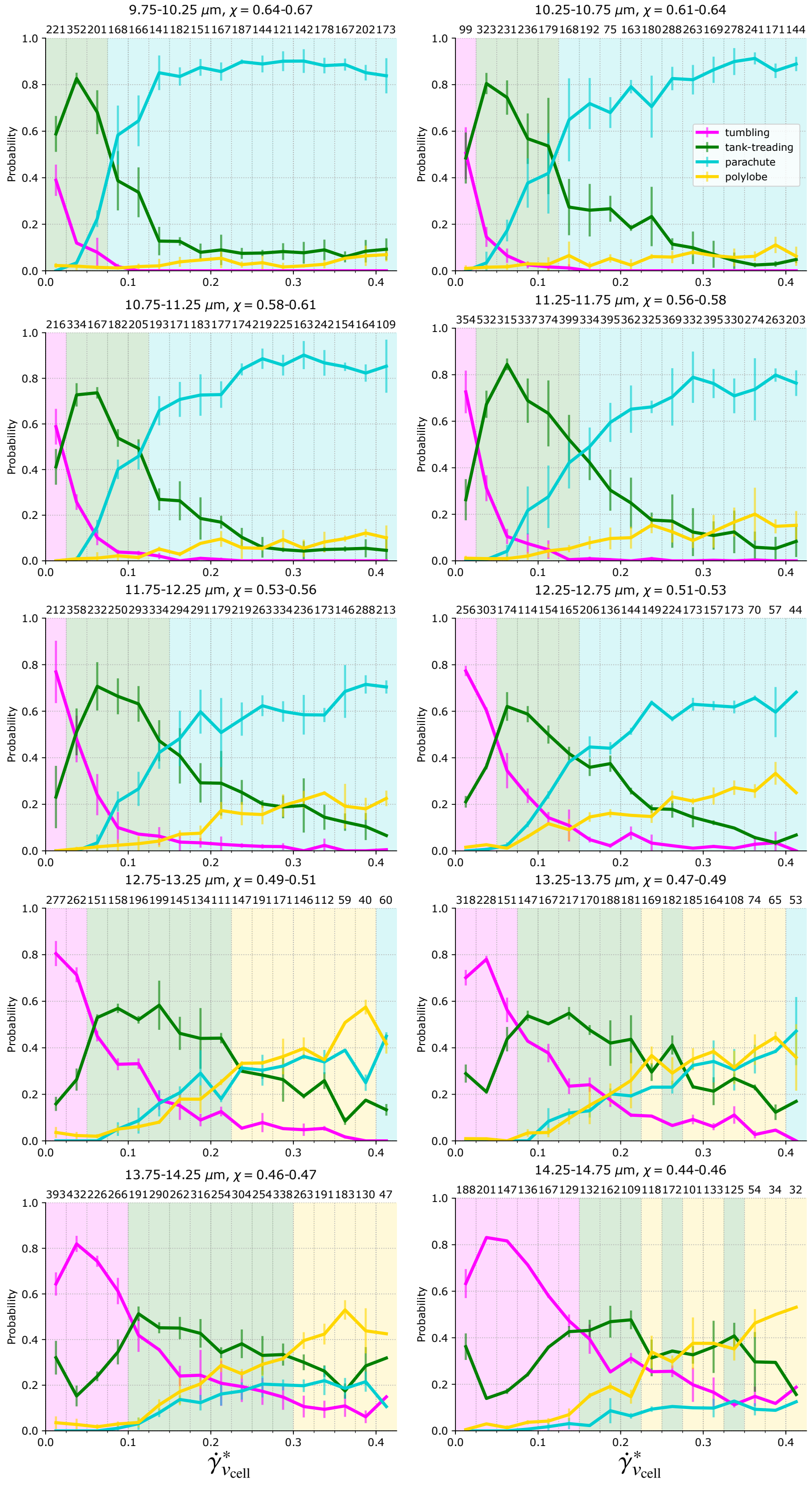


**FIGURE S2.** Shape probability diagrams for all ranges of channel sizes/confinements, as introduced in Fig. 3.

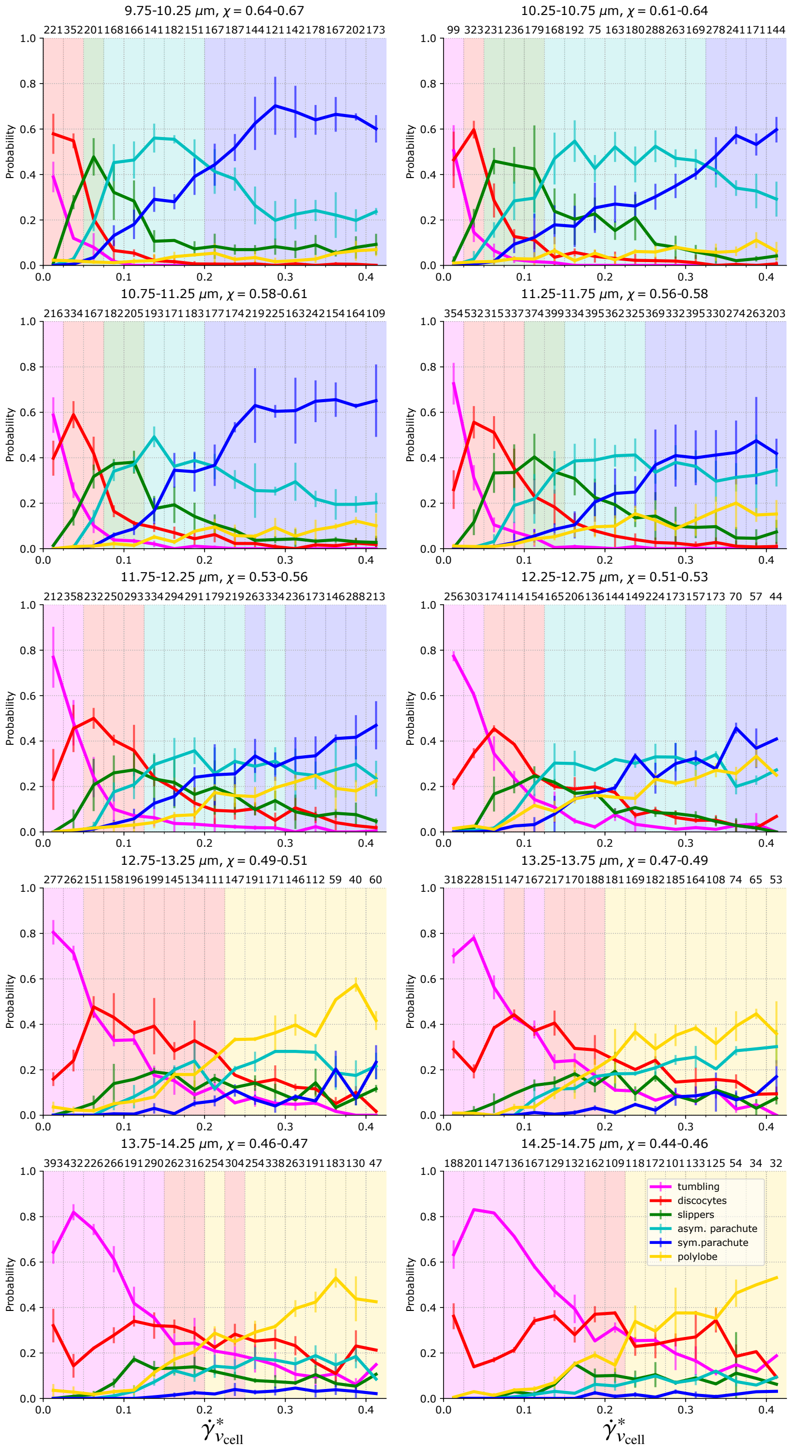


**FIGURE S3.** Shape probability diagrams for all ranges of channel sizes/confinements, further distinguishing between subgroups of tank-treading RBCs and parachutes. Discocytes: subgroup of tank-treading cells that correspond to undeformed RBCs which are not tumbling (see Fig. 2*D*). Slippers: subgroup of tank-treading RBCs deformed into a slipper shape (see Fig. 2*E*). Asymmetrical parachutes: subgroup of parachutes that lack axisymmetry (see Fig 2*F*). Symmetrical parachutes: subgroup of parachutes that show axisymmetry (see Fig 2*G*).

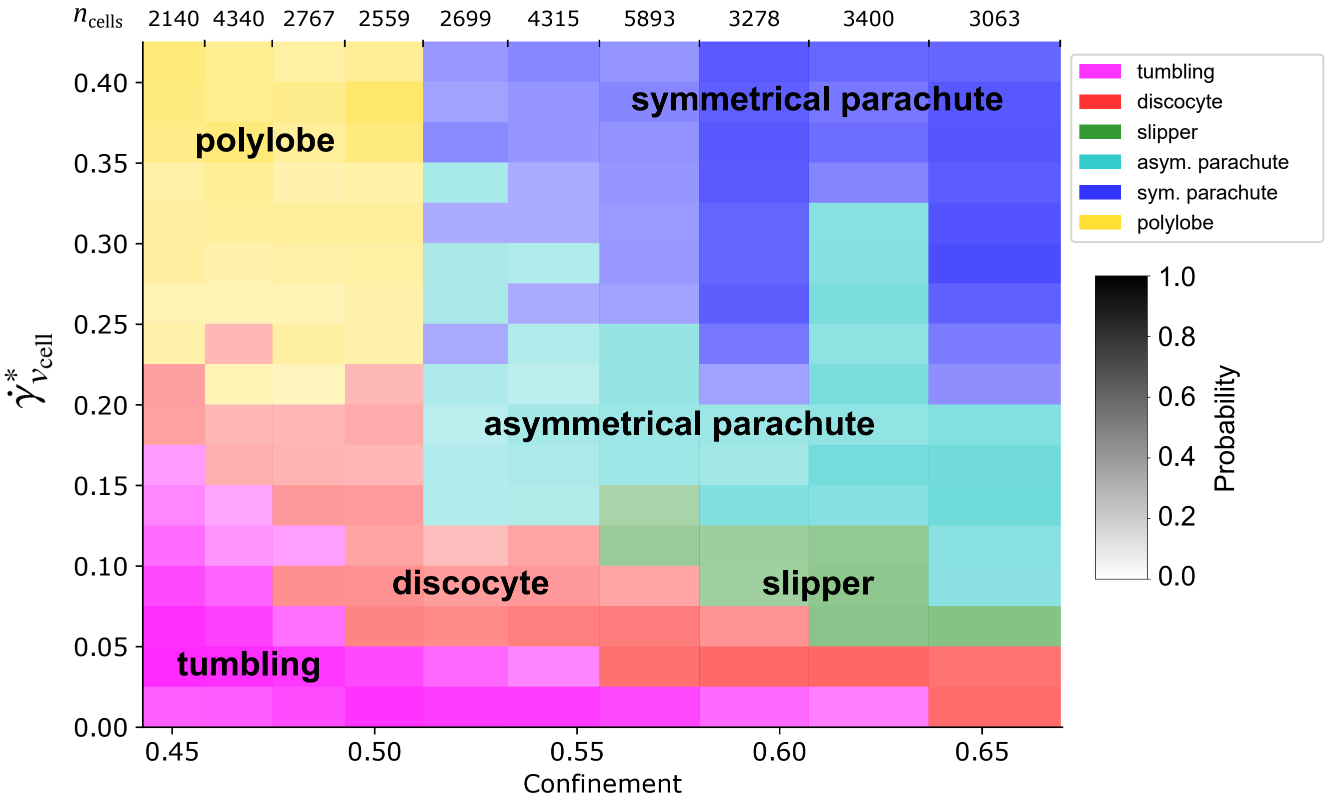


**FIGURE S4.** Phase diagram as in Fig. 3*B* further distinguishing between subgroups of tank-treading cells and parachutes. Discocytes: subgroup of tank-treading RBCs that correspond to undeformed cells which are not tumbling (see Fig. 2*D*). Slippers: subgroup of tank-treading cells deformed into a slipper shape (see Fig. 2*E*). Asymmetrical parachutes: subgroup of parachutes that lack axisymmetry (see Fig 2*F*). Symmetrical parachutes: subgroup of parachutes that show axisymmetry (see Fig 2*G*).

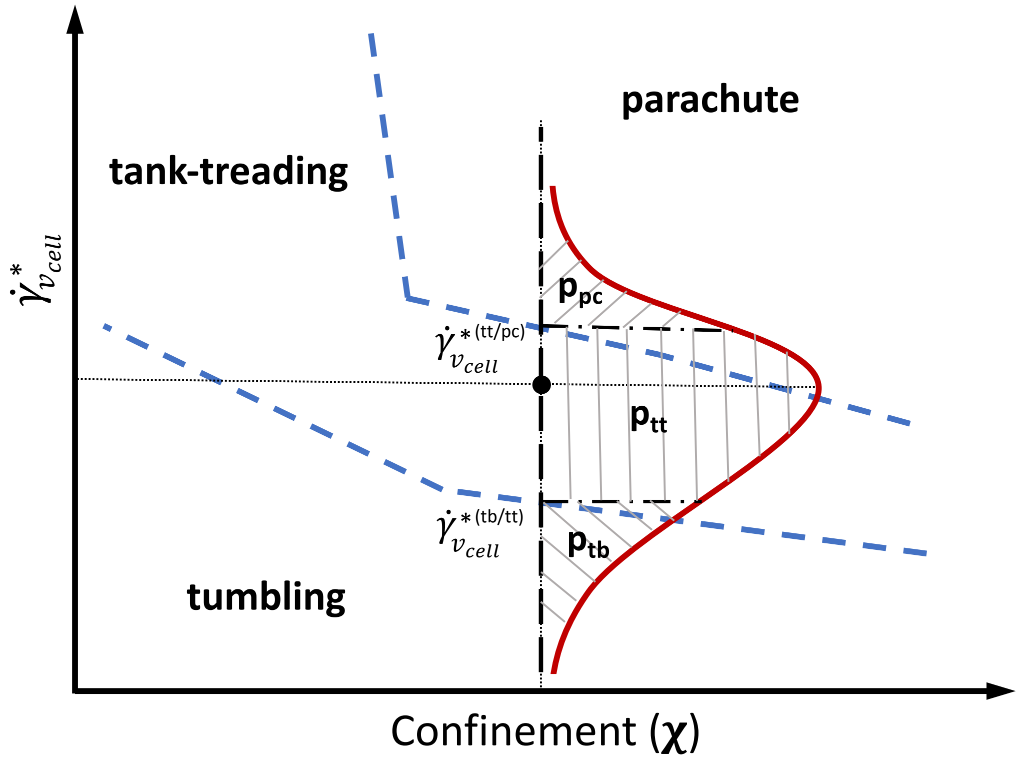


**FIGURE S5.** Schematic superimposition of the variability distribution in shear modulus (see Fig. 4*A*) onto the state diagram in Fig. 3*B*. The shear-modulus distribution is first converted into a $\dot{\gamma}^*_{v_{\text{cell}}}$ distribution (red line), using the relation $\dot{\gamma}^*_{v_{\text{cell}}} \sim 1/\mu_r$. Then, for any selected confinement $\chi$ and $\dot{\gamma}^*_{v_{\text{cell}}}$, the distribution is placed such that its peak coincides with the chosen $\dot{\gamma}^*_{v_{\text{cell}}}$ value. The state boundary points ($\dot{\gamma}^{*(tb/tt)}_{v_{\text{cell}}}$ and $\dot{\gamma}^{*(tt/pc)}_{v_{\text{cell}}}$) between tumbling (tb), tank-treading (tt), and parachute (pc) regions define the integration limits. Integration of the probability distribution over the corresponding ranges ($\dot{\gamma}^*_{v_{\text{cell}}} < \dot{\gamma}^{*(tb/tt)}_{v_{\text{cell}}}$; $\dot{\gamma}^{*(tb/tt)}_{v_{\text{cell}}} < \dot{\gamma}^*_{v_{\text{cell}}} < \dot{\gamma}^{*(tt/pc)}_{v_{\text{cell}}}$; $\dot{\gamma}^{*(tt/pc)}_{v_{\text{cell}}} < \dot{\gamma}^*_{v_{\text{cell}}}$) results in probabilities for observing tumbling ($p_{tb}$), tank-treading ($p_{tt}$), and parachute ($p_{pc}$) states, respectively.

### SUPPLEMENTARY TABLES

| width measurement channel [μm] | width outer branch [μm] |
|---|---|
| 8 | 350 |
| 9 | 350 |
| 10 | 400 |
| 11 | 400 |
| 12 | 400 |
| 13 | 450 |
| 14 | 450 |
| 15 | 450 |

**TABLE S1.** Intended widths of the measurement channel and widths of outer channel branches (see Fig. 2*A*,*B*).

| intended channel height [μm] | spinning frequency [rpm] |
|---|---|
| 10 | 2100 |
| 11 | 2000 |
| 12 | 1900 |
| 13 | 1800 |
| 14 | 1700 |
| 15 | 1600 |

**TABLE S2.** Spinning frequency for the second spinning step of the production of the silicon wafers for each intended channel height.

| Hydraulic diam. [μm] range | Confinement range | Bottom width [μm] | Top width [μm] | Height [μm] | Hydraulic diam. [μm] |
|---|---|---|---|---|---|
| 9.75-10.25 | 0.637-0.670 | 10.71 | 8.98 | 10.38 | 10.07 |
| | | 10.90 | 9.16 | 10.34 | 10.14 |
| | | 11.10 | 9.06 | 10.35 | 10.16 |
| 10.25-10.75 | 0.607-0.637 | 11.08 | 9.74 | 10.29 | 10.32 |
| | | 10.9 | 9.85 | 10.41 | 10.37 |
| | | 11.05 | 9.94 | 10.33 | 10.39 |
| | | 11.00 | 11.21 | 10.23 | 10.64 |
| 10.75-11.25 | 0.580-0.607 | 12.25 | 10.71 | 10.23 | 10.78 |
| | | 11.02 | 9.86 | 11.48 | 10.92 |
| | | 12.00 | 9.86 | 11.48 | 11.14 |
| 11.25-11.75 | 0.556-0.580 | 11.02 | 10.80 | 11.97 | 11.41 |
| | | 11.30 | 10.90 | 12.06 | 11.55 |
| | | 13.60 | 10.83 | 11.16 | 11.57 |
| | | 13.02 | 10.88 | 11.35 | 11.59 |
| | | 11.10 | 11.57 | 11.93 | 11.62 |
| | | 14.40 | 10.97 | 11.03 | 11.65 |
| 11.75-12.25 | 0.533-0.556 | 11.61 | 11.40 | 12.09 | 11.78 |
| | | 14.70 | 11.87 | 11.02 | 11.94 |
| | | 14.80 | 12.14 | 10.92 | 11.97 |
| | | 11.95 | 12.01 | 12.03 | 12.00 |
| | | 11.91 | 11.80 | 12.17 | 12.01 |
| 12.25-12.75 | 0.512-0.533 | 12.99 | 12.80 | 11.89 | 12.37 |
| | | 13.16 | 12.90 | 11.97 | 12.47 |
| 12.75-13.25 | 0.493-0.512 | 14.60 | 11.74 | 13.17 | 13.09 |
| | | 14.42 | 12.12 | 13.27 | 13.22 |
| 13.25-13.75 | 0.475-0.493 | 14.80 | 12.86 | 13.13 | 13.43 |
| | | 15.50 | 12.95 | 13.13 | 13.59 |
| 13.75-14.25 | 0.458-0.493 | 15.98 | 12.91 | 13.41 | 13.81 |
| | | 15.90 | 13.92 | 13.17 | 13.94 |
| | | 16.10 | 14.16 | 13.25 | 14.08 |
| 14.25-14.75 | 0.443-0.458 | 16.50 | 14.10 | 13.74 | 14.42 |
| | | 16.83 | 13.24 | 14.31 | 14.54 |

**TABLE S3.** Channel dimensions based on the schematic shown in Fig. S1. The values were determined on the silicon wafer and on the assembled chip as described in the section Materials and Methods.

### SUPPLEMENTARY MOVIES

1) **Movie S1**: Passage of RBCs from the focusing region to measurement channel (highlighted with red rectangle in Fig. 2).
2) **Movie S2**: Cells passing the serpentine region and detaching from the channel wall.
3) **Movie S3**: Tumbling RBC from simulations for $\chi = 0.47$ at $\dot{\gamma}^*_{v_\mathrm{cell}} = 0.065$ ($v_\mathrm{cell} = 0.74$ mm/s).
4) **Movie S4**: Simulated tank-treading cell for $\chi = 0.47$ at $\dot{\gamma}^*_{v_\mathrm{cell}} = 0.36$ ($v_\mathrm{cell} = 4.1$ mm/s).
5) **Movie S5**: Parachute shape of a RBC from simulations for $\chi = 0.59$ at $\dot{\gamma}^*_{v_\mathrm{cell}} = 0.24$ ($v_\mathrm{cell} = 2.2$ mm/s).
6) **Movie S6**: Channel passage of a tumbling RBC for $\chi = 0.55$ at $\dot{\gamma}^*_{v_\mathrm{cell}} = 0.015$ ($v_\mathrm{cell} = 0.146$ mm/s). The bottom section shows a magnified image that follows the cell during the passage.
7) **Movie S7**: Discocyte with a fixed inclination angle, corresponding to a non-deformed tank-treading cell for $\chi = 0.57$ at $\dot{\gamma}^*_{v_\mathrm{cell}} = 0.103$ ($v_\mathrm{cell} = 0.963$ mm/s).
8) **Movie S8**: Slipper/deformed tank-treading RBC for $\chi = 0.56$ at $\dot{\gamma}^*_{v_\mathrm{cell}} = 0.1$ ($v_\mathrm{cell} = 0.951$ mm/s).
9) **Movie S9**: Asymmetric parachute for $\chi = 0.56$ at $\dot{\gamma}^*_{v_\mathrm{cell}} = 0.233$ ($v_\mathrm{cell} = 2.207$ mm/s).
10) **Movie S10**: Symmetrical parachute for $\chi = 0.56$ at $\dot{\gamma}^*_{v_\mathrm{cell}} = 0.429$ ($v_\mathrm{cell} = 4.066$ mm/s).
11) **Movie S11**: Rigid trilobe/polylobed cell for $\chi = 0.64$ at $\dot{\gamma}^*_{v_\mathrm{cell}} = 0.408$ ($v_\mathrm{cell} = 3.378$ mm/s).
12) **Movie S12**: Shape shifting polylobed cell for $\chi = 0.59$ at $\dot{\gamma}^*_{v_\mathrm{cell}} = 0.345$ ($v_\mathrm{cell} = 3.14$ mm/s).
13) **Movie S13**: Cell changing from asymmetrical parachute to tumbling polylobe for $\chi = 0.6$ at $\dot{\gamma}^*_{v_\mathrm{cell}} = 0.651$ ($v_\mathrm{cell} = 5.788$ mm/s).
14) **Movie S14**: Cell changing from tumbling polylobe to asymmetrical parachute for $\chi = 0.6$ at $\dot{\gamma}^*_{v_\mathrm{cell}} = 0.702$ ($v_\mathrm{cell} = 6.241$ mm/s).
